# The regulatory logic of a dose-dependent developmental fate decision

**DOI:** 10.64898/2026.06.01.729432

**Authors:** Alison H. Araten, Emily K. Ho, Jared E. Toettcher

## Abstract

In canonical developmental patterning, the embryo is exposed to gradients of signaling activators that elicit different cellular responses depending on the activator’s concentration. Recent optogenetic studies of terminal ERK signaling downstream of Torso receptor tyrosine kinase in the early *Drosophila* embryo reveal that even a brief, 5-minute ERK stimulus is sufficient to rescue the development of larval “tail” structures. Here, we reveal components of the molecular network that defines this sensitive developmental fate response. We find that low ERK doses produce sustained *Abdominal-B* (*Abd-B*) expression comparable to that of wild-type embryos. *Abd-B* expression is adjacent to, but non-overlapping with, two other transcriptional repressors: the ERK effector Tailless (Tll) and the gap gene Giant (Gt). Analysis of gene expression patterns in response to optogenetic perturbations suggests that the Tll-dependent repression of *gt* constitutes the sensitive ERK-responsive step: even low *tll* expression leads to potent repression of *gt* in nearby regions, with *Abd-B* expression arising in a stripe between the *tll* and *gt* domains. Our work suggests that the spectrum of phenotypes produced through optogenetic manipulation can be used to define how robust patterning can arise from low doses of inductive signals.

**Highlights:**

- A very low dose of receptor tyrosine kinase signaling in the early embryo induces tail formation many hours later.
- Transient ERK activity results in stable Abd-B expression, beginning as a stripe in nuclear cycle 14.
- Optogenetic ERK inputs induce a spectrum of ectopic phenotypes that reveal mutually exclusive expression of Tailless, Giant, and Abd-B in the posterior.
- Tailless-dependent repression of *giant* is the sensitive ERK-responsive step.

## Introduction

During embryogenesis, developmental signaling patterns give rise to complex spatial organization of cell fates and body structures. Yet the mapping between signaling and fates is not one-to-one: a single signaling pattern can specify multiple fates, with cells adopting different outcomes depending on the signaling dose they receive (Kicheva and Briscoe, 2023). In many cases, it remains unknown how subtle differences in signaling strength can produce precise boundaries between cell fate outcomes. Answering this question is challenging because developmental patterning networks tend to be complex, redundant, and densely interconnected, and because it remains technically challenging to precisely manipulate developmental patterns to systematically interrogate how differences in their dose, duration, or spatial range affect cell fate outcomes.

Terminal patterning at the *Drosophila* embryo’s posterior pole is a canonical example of a single developmental pattern which specifies multiple cell fates. Local binding of the Trunk (Trk) ligand to the Torso (Tor) receptor tyrosine kinase (RTK) initiates a signaling cascade that produces a graded pattern of extracellular signal-regulated kinase (ERK) activity (**Fig 1A**) (Casali and Casanova, 2001; Casanova and Struhl, 1989; Ho et al., 2025; Li, 2005; Lu et al., 1993). Active, doubly-phosphorylated ERK (dpERK) regulates the expression of two transcription factors, Tailless (Tll), which responds sensitively to ERK and is expressed throughout the gradient in a characteristic posterior cap; and Huckebein (Hkb), which is only sensitive to higher ERK doses, leading to a narrower expression domain (Bronner and Jackle, 1991; Jimenez et al., 2000; Smits and Shvartsman, 2020; Weigel et al., 1990). This terminal ERK gradient and its downstream targets are necessary for patterning the embryo’s posterior body structures, which form much later in development after ERK signaling is no longer active (Strecker et al., 1988, 1986).

**Figure 1.**
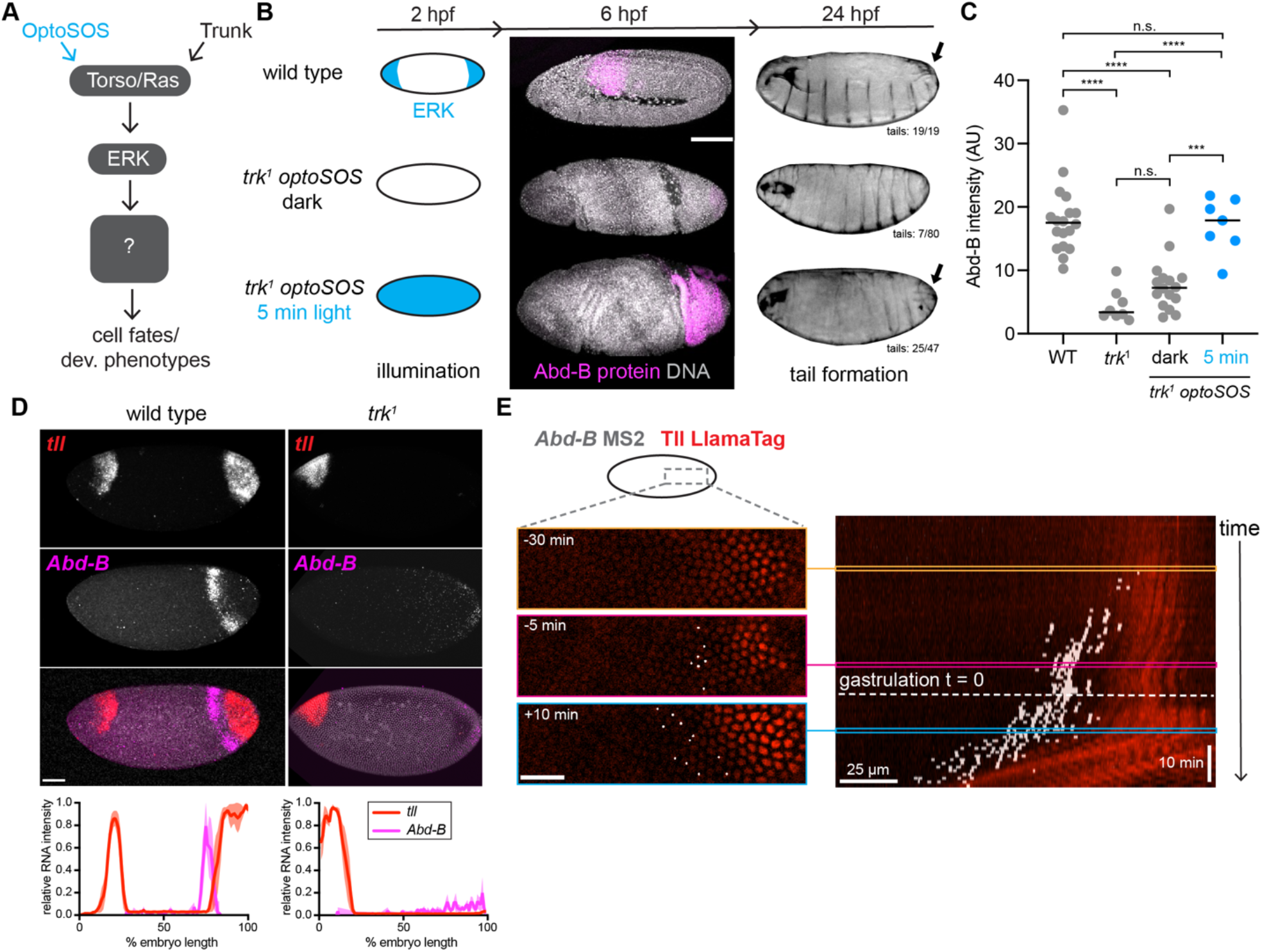
A low dose of ERK signaling is sufficient for tail formation and Abd-B expression. (A) Natural input into the Ras/ERK pathway comes from Trunk binding to Torso RTK, but synthetic, optogenetic input can be achieved using OptoSOS. (B) A short global pulse of ERK activity rescues tail formation and Abd-B expression. Left: Schematic of ERK activity in each genotype and illumination condition (Top: wild-type embryos with endogenous ERK activity. Middle: *trk*^*1*^ *optoSOS* embryos in the dark have no ERK activity. Bottom: *trk*^*1*^ *optoSOS* with a 5-minute pulse of blue light have global ERK activity). Center: Abd-B protein localization at the germband extension stage in a single Z slice. Scale bar 100 μm. Right: Cuticle preps with arrows highlighting posterior spiracles (tail). Proportion indicates number of embryos containing spiracles over total embryos counted. (C) Quantification of Abd-B protein intensity in the tail region of each genotype. Each dot represents 1 embryo. n = 18 (WT), 8 (*trk*^*1*^), 16 (*trk*^*1*^ *optoSOS-* dark), 7 (*trk*^*1*^ *optoSOS-* 5 min light) embryos. ANOVA with Tukey’s post hoc test. Significance defined as *** P<0.001, **** P<0.0001, ns indicates no significance. (D) Maximum intensity projections of *tll* and *Abd-B* RNA in wild-type and *trk*^*1*^ embryos. Scale bar 50 μm. Quantification shows intensity of *tll* and *Abd-B* RNA along the anterior-posterior axis. Line shows mean ± s.e.m. of n = 3 embryos in each condition. Intensity is normalized from 0 to 100 for each gene and within each embryo. (E) Live imaging of *Abd-B* MS2 and Tll LlamaTag in the embryo posterior region indicated by the schematic. Time = 0 is the start of gastrulation movements. Left: Still images show *Abd-B* MS2 foci and Tll LlamaTag expression 30 minutes before, 5 minutes before, and 10 minutes after gastrulation. *Abd-B* MS2 foci have been manually marked with white dots to allow visualization. Right: A kymograph of this region. Gastrulation onset is marked by the dotted line. Colored lines show where each frame falls within the kymographs. Scale bars are 25 μm, 10 min.

In recent years, optogenetic tools have enabled precise manipulation of developmental patterns that can complement traditional genetic analyses (Gao et al., 2026). The OptoSOS system confers this type of manipulation over ERK signaling in the early *Drosophila* embryo (**Fig. 1A**) (Johnson et al., 2017). Varying the dose of blue light tunes the level of ERK activity (Ho et al., 2023; Johnson and Toettcher, 2019), and sequences of light and dark can drive reversible changes in *tll* gene expression on a timescale of minutes (Keenan et al., 2020). When applied in *trk* mutant embryos that lack an endogenous ERK pattern, light is sufficient to drive formation of terminal structures, even fully rescuing development in a proportion of embryos (Johnson et al., 2020). Notably, while complete rescue requires ∼1 h of sustained ERK activity, a stimulus of only 5 min is sufficient for the formation of posterior spiracles or “tails”, anatomical structures that are part of the larval breathing apparatus (Johnson et al., 2020). These data suggest that different target programs respond to ERK doses spanning a ∼10-fold range, and raise the question of how such a brief, weak stimulus can be robustly interpreted to form permanent anatomical structures at the appropriate location.

Here, we combine optogenetic stimuli and imaging to define the key molecular components involved in transducing low doses of RTK signaling in the embryo to larval tail structures. We find that the Hox gene *Abdominal B (Abd-B)* is potently and stably activated in response to very low, transient ERK stimuli, supporting Abd-B induction as a key decision point in the process of tail formation. We identify a network of three transcription factors (TFs) – Tll, Giant (Gt), and Abd-B – whose expression patterns are mutually exclusive in the posterior domains of wildtype and optogenetically stimulated embryos. High levels of Tll repress *Abd-B* whereas low levels of Tll serve to activate *Abd-B*, either directly or by inhibiting *gt* expression. Our data suggest that Tll repression of *gt* is the exquisitely sensitive step that amplifies a weak stimulus into an all-or-none response, promoting establishment of a narrow stripe of Abd-B in a domain that lacks both Tll and Gt expression.

## Results

### A low dose of ERK signaling is sufficient for full rescue of Abd-B expression

We first set out to identify the key molecular players that interpret low RTK signaling doses to form posterior spiracles (tail structures). Given that a low, transient dose of ERK activity in the early embryo induces tail structures ∼20 h later, we reasoned that a “tail regulator” might be induced in the early embryo by ERK and persist through development. The Hox genes are compelling candidates for this role; their expression is induced early by gap and pair rule genes and then maintained over developmental time to define the identity of each segment (Maeda and Karch, 2006). Posterior spiracles form in the domain of the most posterior Hox gene, Abdominal B (Abd-B). Abd-B is known to be necessary and sufficient for the formation of posterior spiracles (Castelli-Gair et al., 1994; Hombría et al., 2009; Hu and Castelli-Gair, 1999), and Abd-B has also been genetically linked to expression of the ERK target *tll* (Reinitz and Levine, 1990). Thus, we hypothesized that induction of Abd-B expression might provide the link between a low ERK pulse and tail formation.

In the wild-type embryo, terminal RTK signaling at 2 h post fertilization (hpf) leads to a robust domain of posterior Abd-B protein by at least 6 hpf, during the germband elongation stage (**Fig 1B**; top). In these embryos, tail structures form normally and can be detected in the cuticle at the time of hatching (**Fig 1B**; top). In dark-incubated *trk*^*1*^ *optoSOS* embryos that lack both endogenous and light-induced terminal ERK signaling, neither Abd-B protein expression nor tail structures are detected, confirming the genetic link between ERK signaling, Abd-B, and tails (**Fig 1B**; middle). We note that in these embryos that lack proper terminal patterning, failed posterior midgut formation and invagination cause a twisting gastrulation phenotype that prevents germ-band elongation, leading to a different position of Abd-B expressing cells within the embryo (Johnson et al., 2020; Smits et al., 2023).

In contrast, *trk*^*1*^ *optoSOS* embryos stimulated with a 5-min pulse of saturating 450 nm blue light (0.617 mW/cm^2^) exhibited tail formation in ∼50% of embryos (Johnson et al., 2020), and all embryos showed a robust domain of Abd-B protein expression at the posterior of the twisted, 6 h embryos (**Fig 1B**, bottom). Quantification of Abd-B levels in each condition showed that the 5-min light pulse induces a significant increase in posterior Abd-B expression compared to embryos kept in the dark, resulting in expression comparable to that observed in wild-type embryos (**Fig 1C**). Quantification also revealed substantial heterogeneity in Abd-B expression across dark-incubated *trk*^*1*^ *optoSOS* embryos (**Fig 1C**), suggesting that this optogenetic system may exhibit some “leaky” ERK activity in the dark that is sufficient to partially activate Abd-B (**Fig 1B**; middle). Nevertheless, our results demonstrate that the low dose of ERK signaling that is sufficient for tail formation is also sufficient to induce robust, sustained Abd-B expression, consistent with Abd-B’s expected role linking posterior ERK signaling to downstream tail formation.

### A narrow stripe of *Abd-B* expression emerges in late NC14

Classic genetic studies suggest that the ERK target gene Tll is a direct or indirect activator of Abd-B, as Abd-B expression is absent in *tll* mutant embryos (Harding and Levine, 1988; Reinitz and Levine, 1990). Therefore, we sought to examine the relationship between Tll and Abd-B expression, hypothesizing that the timing and/or position of Abd-B expression relative to Tll might provide insights into its sensitivity to low doses. For this purpose, we reasoned that tools for detecting *Abd-B* RNA would provide a more proximal readout of Abd-B’s induction.

We used hybridization chain reaction (HCR) to detect *Abd-B* transcripts in fixed embryos. *Abd-B* is expressed in two isoforms, a morphogenic isoform (*Abd-Bm*) and a regulatory isoform (*Abd-Br*), where the morphogenic isoform is thought to correspond more closely to functional responses including posterior spiracle formation (Kuziora, 1993; Kuziora and McGinnis, 1988). To disentangle these responses, we developed HCR probes to measure either total *Abd-B* or the regulatory *Abd-Br* isoform and applied each probe to wild-type embryos (**Supplementary Fig. 1A**). We found that the total *Abd-B* probe corresponded closely to the expected pattern of the morphogenic isoform, whereas a regulatory isoform-only probe revealed a weaker and more posterior-shifted pattern (**Supplementary Fig. 1B**). Thus, we used the total *Abd-B* probe for the remainder of the paper.

We combined our *Abd-B* probe with a probe for *tll* to examine both genes in nuclear cycle 14 (NC14), a period where terminal ERK signaling induces target gene expression. We detected a stripe of *Abd-B* transcripts in late NC14 embryos at about 75% embryo length (**Fig 1D**; left), consistent with prior work (Harding and Levine, 1988). At this stage, *tll* expression forms a posterior cap. Quantification of *Abd-B* and *tll* together showed that the *Abd-B* stripe sits at the anterior edge of the posterior *tll* expression domain, adjacent to but not overlapping the domain of detectable *tll* expression. At this position, Tll levels are presumably very low, consistent with induction of *Abd-B* by low ERK. Additionally, in *trk*^*1*^ mutants, *Abd-B* and posterior *tll* expression are both absent, supporting the requirement of ERK activity and Tll for expression of *Abd-B* (**Fig 1D**; right).

We then asked whether this stripe of *Abd-B* forms immediately as a stripe or whether it is refined from an initial broader pattern of expression, as this distinction is important for informing possible models of regulation (Birnie et al., 2023; Ho et al., 2023; Keenan et al., 2022; Zhao et al., 2023). We developed complementary live-embryo MS2 biosensors to monitor the dynamics of Tll-to-*Abd-B* regulation with fine spatiotemporal resolution. We again produced two isoform-specific *Abd-B* MS2 probes to detect total *Abd-B* or the regulatory isoform, but focused our investigation on the total *Abd-B* MS2 allele (**Supplementary Fig. 1C, D**). We produced embryos that co-expressed this MS2-tagged *Abd-B* allele with an existing Tll LlamaTag for real-time protein visualization of the transcription factor (Ho et al., 2023) (**Fig. 1E**). Live imaging experiments revealed that Tll protein accumulates in posterior nuclei approximately 30 min prior to detectable transcription of *Abd-B* (**Fig. 1E**, right). They also demonstrated that *Abd-B* is always expressed as a narrow stripe without substantial overlap with the domain of Tll protein expression. This behavior is distinct from other stripe-forming genes whose initially broad expression refines over time, including *brachyenteron* and *even skipped* (Bothma et al., 2014; Ho et al., 2023; Keenan et al., 2022). These data suggest the factors which specify the domain of *Abd-B* expression are already present at the time that its expression first occurs, rather than being gradually adjusted and refined over time. They also prompt the question of how Tll regulation establishes an Abd-B stripe at this unique position.

### Optogenetic stimuli produce shifted, non-overlapping *tll* and *Abd-B* domains

What is the regulatory logic by which Tll influences *Abd-B* expression? We reasoned that one way to gain insight into this question would be to perturb the domain of *tll* expression and monitor resulting changes in the *Abd-B* pattern (**Fig. 2A**). We turned to the OptoSOS optogenetic system, as our prior work demonstrated that the spatial domain of *tll* expression could be expanded anteriorly by a global increase in light-induced ERK activity (**Fig. 2B**) (Johnson and Toettcher, 2019).

**Figure 2.**
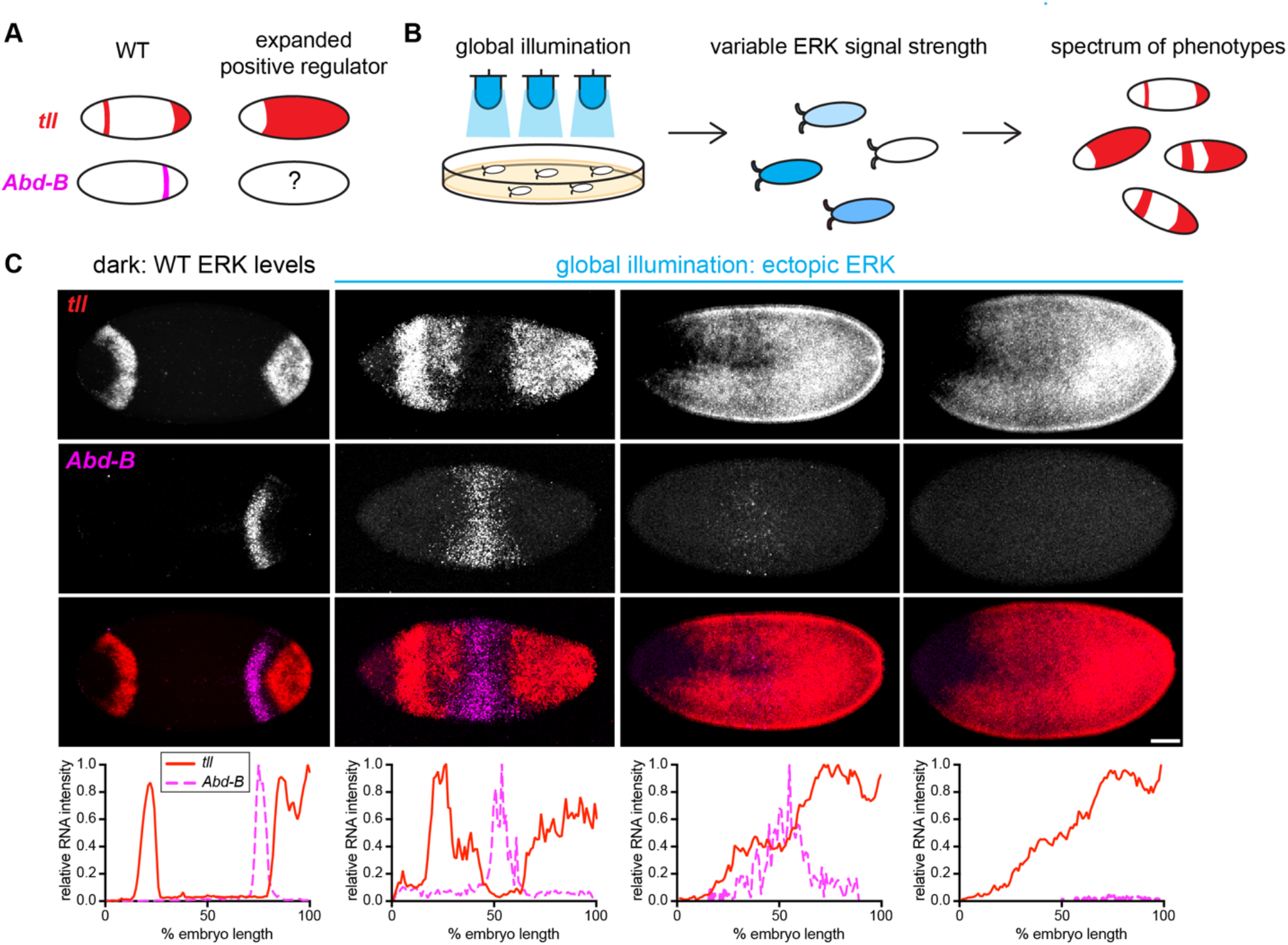
Optogenetic ERK stimuli shift *tll* and *Abd-B* expression domains. (A) In wild-type embryos, the *Abd-B* stripe forms just anterior to the posterior *tll* domain. If expression of *tll* expands, what pattern of *Abd-B* will result? (B) Because of variability in *optoSOS* expression level, bulk illumination of *optoSOS* embryos results in different levels of ERK signaling and therefore a spectrum of *tll* expansion phenotypes. (C) Maximum projection images show a spectrum of *tll* expansion phenotypes and the corresponding *Abd-B* expression patterns in NC14. The wild-type pattern is found in *optoSOS* embryos kept in the dark. When exposed to 2 hrs of light, a spectrum of expansion patterns results. Scale bar 50 μm. Graphs below plot the normalized intensity of each gene across the anterior-posterior axis for the embryo pictured. Intensity is normalized from 0 to 100 for each gene and within each embryo.

We first stained for *tll* and *Abd-B* transcripts in dark-incubated OptoSOS embryos, confirming that these embryos exhibited wild-type patterns of both genes: a narrow stripe of *Abd-B* expression just outside the boundary of the posterior *tll* cap (**Fig. 2C**, first column). We next applied global optogenetic stimuli to perturb ERK activity and monitor *tll* and *Abd-B* responses. Embryos were placed under a lower intensity, 0.07 mW/cm^2^ blue light for 2 h to achieve a spectrum of intermediate phenotypes that likely depends on each embryo’s OptoSOS expression level (Ho et al., 2023). This variability proved extremely useful, as we were able to observe a spectrum of perturbations to gene expression patterning, allowing us to gain a more thorough understanding of the regulatory relationship between *tll* and *Abd-B* (**Fig 2B**).

Previous measurements of *tll* expansion in response to global optogenetic ERK inputs have shown that at increasing ERK input levels, the posterior and anterior domains of *tll* progressively expand towards the center of the embryo (Johnson and Toettcher, 2019), likely due to repressive interactions with Bicoid and the gap gene network that render the anterior-most and central regions of the embryo resistant to ERK-induced *tll* expression. Our globally illuminated embryos stained for *tll* and *Abd-B* transcripts displayed gene expression patterns consistent with these previous measurements, exhibiting a range of different *tll* expansion phenotypes (**Fig 2C**). In embryos that display comparatively weak phenotypes, the anterior and posterior domains of *tll* expression were broader than in wildtype embryos (**Fig 2C**, second column). In these “partial *tll* expansion” embryos, *Abd-B* expression was similarly shifted into regions where *tll* expression is not detected. In a second class of embryos, *tll* was expressed throughout the embryo, except for the head region (**Fig 2C**, third and fourth column). In some of these “total *tll* expansion” embryos we observed a faint stripe of *Abd-B* expression overlapping with the region of lowest *tll* expression in the center of the embryo (**Fig 2C**, third column). In others, we observed no *Abd-B* (**Fig 2C**, fourth column). This spectrum of phenotypes suggests a complex regulatory relationship between *tll* and *Abd-B*: ERK-induced *tll* is required for *Abd-B* expression and yet high *tll* expression levels are incompatible with *Abd-B* expression. Our results also demonstrate that optogenetic tools are well-suited to generating spectra of phenotypes, akin to allelic series that have been so useful in dissecting genetic pathways (Diaz et al., 1996; Strecker et al., 1988).

### A short optogenetic pulse of ERK signaling rescues the *Abd-B* stripe

Our data so far are consistent with a model where *Abd-B* expression is promoted by very low levels of Tll protein at the boundary of the *tll* expression domain, but repressed within this domain. We asked whether this model might also explain tail formation in a different scenario: the formation of tail structures in *trk*^*1*^ *optoSOS* embryos upon global illumination with low-dose light stimuli. Remarkably, these embryos form tails in the correct position despite the fact that the ERK-activating light input is delivered uniformly to the entire embryo. How do *tll* and *Abd-B* respond to brief global illumination, and are these patterns consistent with our inferred model (**Fig. 3A**)?

**Figure 3.**
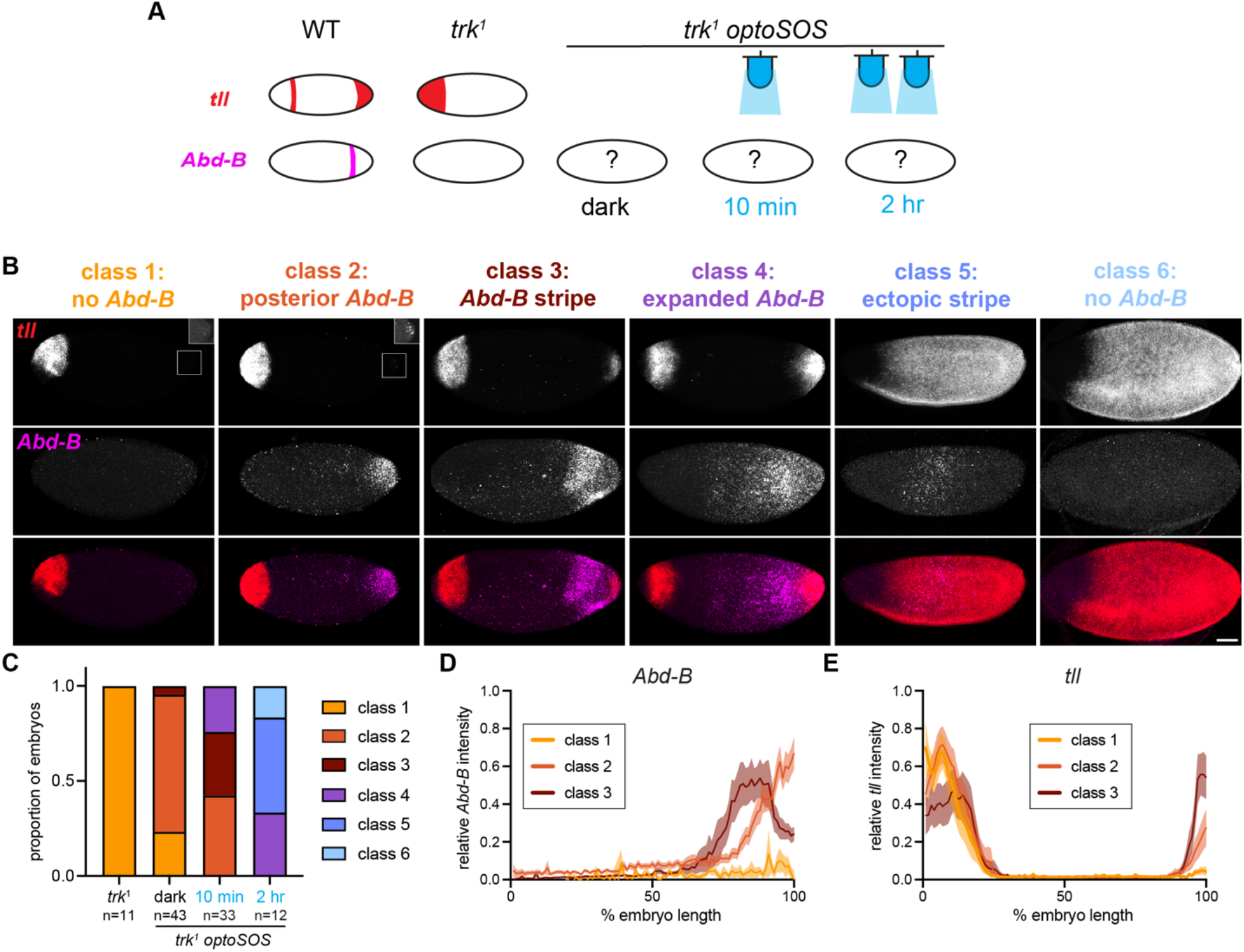
Optogenetic expansion of *tll* and *Abd-B* expression in *trk* mutant embryos. (A) The posterior domains of *tll* and *Abd-B* are lost in t*rk*^*1*^ mutants lacking ERK signaling. ERK signaling can be restored at differing levels by illumination of *trk*^*1*^ *optoSOS* embryos. (B) Four groups of embryos were imaged in NC14: *trk*^*1*^ mutants, *trk*^*1*^ *optoSOS* – dark, *trk*^*1*^ *optoSOS* – 10 min illumination at 0.617 mW/cm2, and *trk*^*1*^ *optoSOS* – 2 h illumination at 0.077 mW/cm2. All embryos were sorted into classes based on the *Abd-B* expression phenotype. Class 1 embryos have no or barely detectable *Abd-B*. Class 2 embryos have strong *Abd-B* that is localized to only the most posterior. Class 3 embryos have a distinct stripe of *Abd-B*. In class 4 embryos, the domain of *Abd-B* extending from the posterior is large. In Class 5, *Abd-B* is only expressed in the center of the embryo, not at the posterior. In Class 6, there is no *Abd-B* expression (but strong *tll* expression throughout the embryo). Insets for class 1 and class 2 show posterior *tll* at a higher brightness. Images show representative embryos in each class. Scale bar 50 μm. (C) For each condition, the proportion of embryos sorted into each class is shown. The number of embryos for each condition is indicated on the graph. (D, E) Quantification of *Abd-B* (D) and *tll* (E) intensity across the anterior-posterior axis for all embryos in classes 1-3. Line shows mean ± s.e.m. of n = 8 (class 1), 13 (class 2), 9 (class 3) embryos in each condition. Intensity is normalized from 0 to 100 for each gene and within each embryo.

We incubated *trk*^*1*^ *optoSOS* embryos in three different illumination conditions (dark, 10 min light, 2 h light) and compared the resulting *tll* and *Abd-B* expression patterns to those of *trk*^*1*^ embryos. We classified each embryo according to the pattern of *Abd-B* expression we observed (**Fig 3B**) and then, for each stimulation condition, determined the proportion of embryos in each class (**Fig 3C**). Consistent with an absence of terminal ERK activity, all *trk*^*1*^ embryos lacked posterior *tll* and *Abd-B* expression, which we define as a “class 1” phenotype (**Fig 3B** column 1, **3C**). In *trk*^*1*^ *optoSOS* embryos kept in the dark we again noted some leakiness with the OptoSOS construct, leading to a mix of class 1 and class 2 embryos. These class 2 embryos exhibited very low levels of *tll* expression and robust *Abd-B* expression in a posterior domain, consistent with low *tll* levels favoring *Abd-B* expression (**Fig 3B** columns 1-2, **3C**). For embryos stimulated with a transient 10-min pulse of bright light (0.617 mW/cm^2^), we observed some class 2 embryos, but the majority of embryos showed either a stripe of *Abd-B* expression (class 3) or a large posterior domain of expression (class 4) (**Fig 3B** columns 2-4, **3C**). These two classes correlated with increasing domains of *tll* expression at the posterior. For embryos stimulated with 2 hours of global illumination, we observed a mix of class 4 embryos with the two fully expanded tailless phenotypes we previously observed in wild-type embryos: full *tll* expansion with a central Abd-B stripe (class 5) or no Abd-B expression (class 6) **(Fig. 3B**, columns 4-6, **3C**).

Quantifying gene expression across this spectrum of phenotypes (**Fig. 3D-E**) supports a model where proper *Abd-B* patterning is established by a complex dose dependence on ERK activity and *tll* expression. In the complete absence of ERK activity (class 1), neither *tll* nor *Abd-B* expression is induced. When ERK activity is extremely low (class 2), weak *tll* expression is detected along with robust expression of *Abd-B*. As ERK inputs increase further (class 3), increasing posterior *tll* levels inhibit *Abd-B* expression, leading to formation of a gene expression stripe that may be responsible for rescued tail formation.

### Tll represses *gt* expression with exquisite sensitivity

Our investigation thus far suggests that Tll both activates and inhibits *Abd-B* expression in a dose-dependent manner (**Fig. 4A**). Inhibition of *Abd-B* to set the posterior boundary is consistent with Tll’s well-characterized role as a direct transcriptional repressor (Masuda et al., 2024; Morán and Jiménez, 2006). However, how *Abd-B* can be activated by Tll with even greater sensitivity to set the anterior boundary is less clear (**Fig. 4A**). A compelling model is that an additional target gene, likely a transcriptional repressor, is repressed by Tll with exceptionally high sensitivity, clearing a zone for *Abd-B* expression. A candidate gene that is consistent with this model would be expected to be expressed just anterior to the *Abd-B* domain boundary and should be sensitive to perturbations that affect the spatial pattern of *tll* expression.

**Figure 4.**
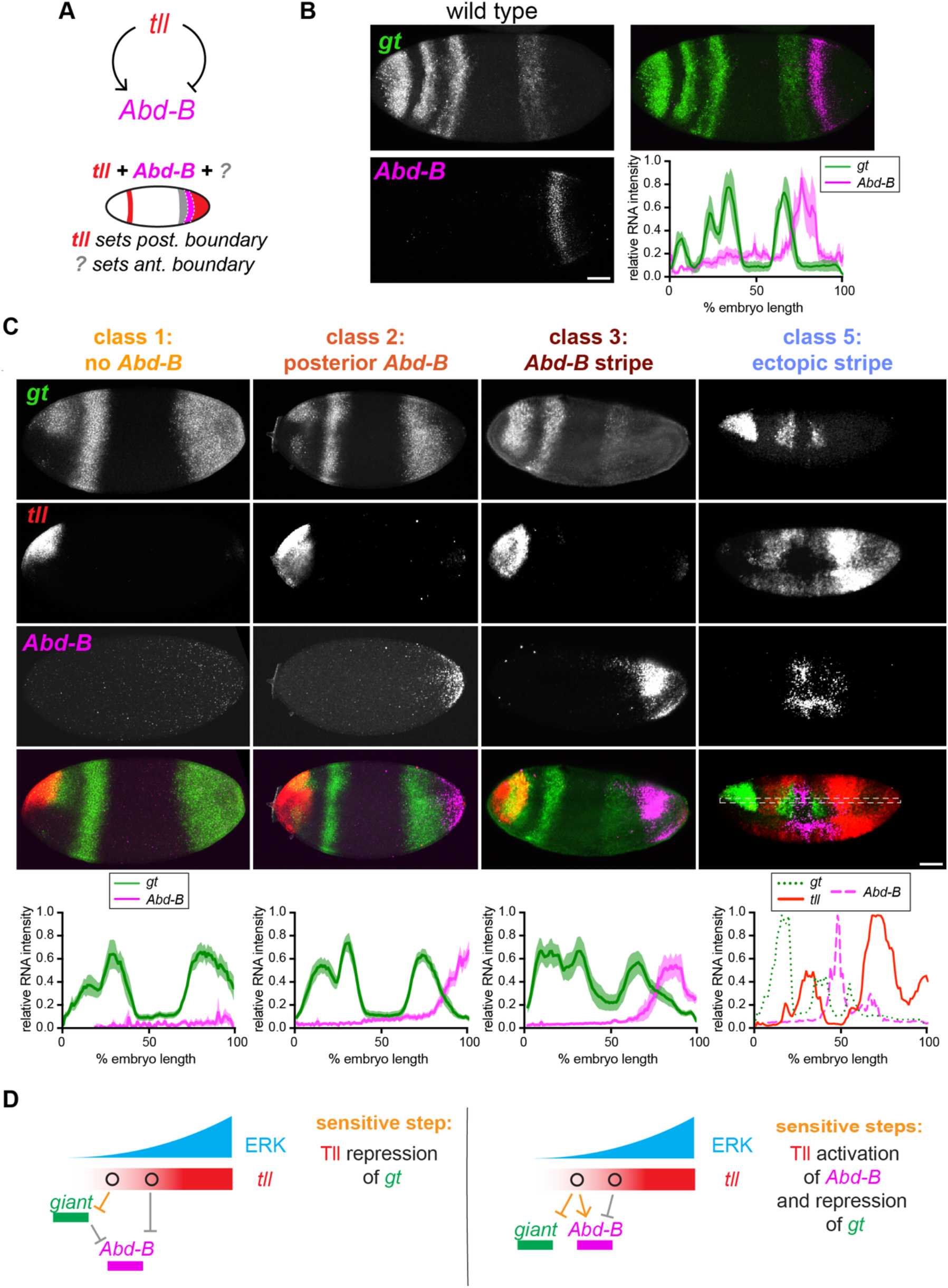
Expression of *gt* and *Abd-B* are mutually exclusive. (A) Tll is a positive and negative regulator of *Abd-B* expression. Tll sets the posterior boundary of Abd-B but which factor sets the anterior boundary? (B) Maximum intensity projection demonstrates the wild-type pattern of *giant* and *Abd-B* in a late NC14 embryos. Embryo shown is an *optoSOS* embryo kept in the dark. Quantification shows *gt* and *Abd-B* intensity across the anterior-posterior axis. Line shows mean ± s.e.m. of n = 3 embryos. (C) Representative embryo in classes 1, 2, 3, and 5 showing the *gt, tll*, and *Abd-B* expression patterns. Quantification of *gt* and *Abd-B* intensity across the anterior-posterior axis for all embryos in classes 1-3 and the individual embryo shown for class 5 (with dotted line indicating region of quantification). Line shows mean ± s.e.m. of n = 8 (class 1), 13 (class 2), 9 (class 3) embryos in each condition. Intensity is normalized from 0 to 100 for each gene and within each embryo. Abd-B quantifications are the same as in Figure 3D. (D) Two possible models of Abd-B regulation by ERK signaling. In both models, the posterior boundary of Abd-B is set by repression by Tll. In model 1 (left), the sensitive step (orange arrow) is repression of *gt* by Tll which relieves repression of *Abd-B*. In model 2 (right), the sensitive step is that low levels of Tll directly activate *Abd-B*.

The other members of the gap gene network, *Kruppel (Kr), knirps (kni), hunchback (hb)* and *giant (gt)*, are compelling candidates for this role, given their mutually repressive relationships with Tll as well as known genetic interactions with Abd-B (Casares and Sanchez-Herrero, 1995; Eldon and Pirrotta, 1991; Harding and Levine, 1988; Reinitz and Levine, 1990). The *Abd-B* stripe expands anteriorly in *kni* mutant embryos, and both *gt* and *Kr* mutant embryos show ectopic *Abd-B* expression. Examining the wildtype expression patterns of each gap gene in relation to *tll* and *Abd-B* expression, it is apparent that *gt* and *Abd-B* expression are most closely opposed (**Supplementary Fig. 2A-D**). In early NC14, *gt* is expressed in a broad posterior region directly adjoining the boundary of *tll* expression. By late NC14, a spatial gap between *gt* and *tll* gives way to the *Abd-B* stripe (**Fig 4B, Supplementary Fig 2A**). In contrast, the domain of *hb* expression overlaps with *Abd-B*, inconsistent with a putative role as a direct repressor (**Supplementary Fig 2B**). The domains of *kni* and *Kr* do not align with the Abd-B boundary (**Supplementary Fig 2C,D**). Based on these data, *gt* appeared to be a top candidate for mediating Tll-dependent de-repression of *Abd-B*.

We again used optogenetic perturbations in *trk*^*1*^ *optoSOS* embryos to gain insight into the interplay between *tll, gt*, and *Abd-B* patterning. In class 1 *trk*^*1*^ or dark *trk*^*1*^ *optoSOS* embryos which completely lack *tll* and *Abd-B* expression, the domain of *gt* expression extends all the way to the posterior of the embryo (**Fig. 4C** column 1). In contrast, class 2 embryos, which are characterized by very low posterior *tll* expression and high posterior *Abd-B*, showed a consistent retraction of *gt* expression from the posterior pole **(Fig. 4C**, column 2). The posterior-most edge of the *gt* stripe consistently bordered but did not overlap the anterior edge of the *Abd-B* domain. Further, in class 3 embryos, which are characterized by an *Abd-B* stripe, we observed a *gt* pattern resembling that of wild-type embryos: *gt* was expressed in a clear stripe, just anterior to the *Abd-B* domain **(Fig. 4C**, column 3). In these class 3 embryos, the *Abd-B* stripe sits between *gt* and *tll*, just as in wild-type embryos. These results suggest that *gt* is extremely sensitive to low level ERK inputs; very weak expression of *tll* is sufficient to push *gt* from the posterior domain.

Overall, we found that *gt, tll* and *Abd-B* are expressed in mutually exclusive domains, even when the expression domains of all three genes are substantially perturbed. An extreme example of this mutual exclusivity can be found in a class 5 embryos with partially expanded *tll* expression and irregular, ectopic regions of *Abd-B* and *gt* expression (**Fig. 4C**, right column). Despite considerable deviations from their usual expression patterns, all three genes are expressed in adjacent but non-overlapping patchy domains. Taken together, these data establish a model where the domain of *Abd-B* expression is established by exquisitely sensitive repression of *gt* by Tll downstream of terminal ERK signaling. It is likely that this repression represents the sensitive step that enables a brief optogenetic pulse to rescue tail formation in *trk*^*1*^ *optoSOS* embryos.

## Discussion

Here, we combined optogenetic perturbations with gene expression assays to dissect the molecular control over an extremely sensitive developmental decision: formation of posterior tail structures. We show that anatomical tail formation is preceded, many hours earlier, by a narrow stripe of *Abd-B* expression whose expression is induced by only low level ERK signaling. We identify Tll as a key regulator of *Abd-B* that transmits information about ERK dose to the position of *Abd-B* within the embryo. Finally, we suggest that one key function of Tll is to prevent expression of the gap gene *gt* within the *Abd-B* expression domain, and we show that Tll repression of *gt* is a key sensitive step in this network.

These data are broadly consistent with two putative models for Abd-B stripe formation (**Fig. 4D**). One model is that *gt* blocks *Abd-B* expression in the absence of ERK stimuli, and Tll indirectly activates *Abd-B* through repression of *gt* (**Fig. 4D** - left). This model is consistent with Tll’s characterization as an exclusive repressor (Masuda et al., 2024; Morán and Jiménez, 2006). However, given that *gt* mutant embryos do not show an anterior expansion of the *Abd-B* stripe, it is likely that Gt acts redundantly with other gap genes such as Kni and Kr to confine *Abd-B* expression to its narrow stripe region (Harding and Levine, 1988; Reinitz and Levine, 1990), or that an as-yet-undefined direct transcriptional activator of *Abd-B* expression also plays an important role in defining its expression domain.

An alternate model is that Tll may have an additional, uncharacterized role as a direct transcriptional activator for *Abd-B* (**Fig. 4D** - right). If low levels of Tll directly activate *Abd-B* expression, the position and width of the *Abd-B* stripe would always be defined solely by the Tll gradient. This could neatly explain why the posterior stripe of *Abd-B* remains narrow in *gt* mutant embryos. A recent optogenetic study of Bicoid-dependent gap gene regulation suggests that this canonical transcriptional activator can act as a repressor of *kni* at low expression levels (Singh et al., 2022), revealing that even classic developmental transcription factors can play unexpected, concentration-dependent regulatory roles. An exciting opportunity for future work will be to distinguish between these models by considering the sensitivities of Tll and Gt transcription factor binding interactions in response to varied ERK inputs, potentially taking advantage of optogenetics to induce these sensitive signaling states (Keenan et al., 2020; Pagella et al., 2023).

Altogether, our data provide molecular clues for how a very low dose of OptoSOS activity can be robustly detected by the embryo and interpreted to produce a permanent anatomical structure (Johnson et al., 2020). We show that a 5-10 min pulse of ERK activity is sufficient to induce *tll* expression, inhibit *gt* expression, and produce a stable stripe of *Abd-B* expression. It was previously shown that transient optogenetic ERK stimulation induces a correspondingly transient pulse of *tll* expression (Keenan et al., 2020), indicating that *Abd-B* converts a transient stimulus into a sustained response. An exciting next step is to seek a systems level understanding of the mechanisms that convert these transient inputs into persistent patterns of gene expression, such as epigenetic changes, autoregulation, and positive feedback (Bowman et al., 2014; Maeda and Karch, 2006; Moniot-Perron et al., 2023; Zhao et al., 2023). For example, simultaneous induction of *hb* may help maintain a domain favorable for *Abd-B* expression through sustained repression of posterior *gt* (Casanova, 1990; Kraut and Levine, 1991; Reinitz and Levine, 1990).

Importantly, our findings do not necessitate that ERK activity occur in a pulse (i.e. Abd-B is not a true “pulse detector”): even 2 h of continuous ERK stimulus produced patchy, non-overlapping domains of *Abd-B, tll*, and *gt* expression in some embryos. However, a pulse of ERK appears to be a convenient means for transmitting very low ERK inputs that produce a transient pulse of *tll* expression and thus very low levels of Tll protein. In wild-type embryos, it is notable that the *Abd-B* stripe forms in a region where ERK activity may only be transiently induced. Previous studies have shown that Torso RTK activity, ERK activity and *tll* expression retract towards the posterior over time, suggesting that the wild-type *Abd-B* stripe may similarly receive a transient pulse of ERK (Clark et al., 2022; Coppey et al., 2008; Ho et al., 2025).

Finally, this work highlights a new strategy for using optogenetics to interrogate developmental networks. In many use cases, variability in optogenetic tool expression level presents a challenge because identical inputs do not induce the same response across multiple embryos (Gao et al., 2026). Here, we show that this phenotypic variability can also be exploited to measure changes in gene expression patterning across of spectrum of stimulus strengths. We hope that this and other strategies for applying developmental optogenetics will uncover new insights from even classical developmental model systems.

## Acknowledgements

We thank Eric Wieschaus, Hernan Garcia, and Mike Levine for the kind gift of flies and plasmids used in this study. We thank Philippe Batut for advice on MS2 reporter generation and Mike Levine, Paul Schedl, Jodi Schottenfeld-Roames and all members of the Toettcher lab for helpful discussions. Confocal imaging was done in the Princeton Confocal Imaging Facility, a Nikon Center of Excellence, with assistance from Gary Laevsky and Sha Wang. Stocks obtained from the Bloomington Drosophila Stock Center (NIH P40OD018537) were used in this study. We thank QG Bu for assistance with stock maintenance. This work was supported by the Pitts ‘63 Senior Thesis Fund (A.H.A.), the National Institutes of Health grants F32GM148016 (E.K.H.); U01DK127429, R01GM144362, and R35GM164185 (J.E.T.).

## Author Contributions

**Alison H. Araten:** Conceptualization, Methodology, Investigation, Formal analysis, Writing-Original Draft, Writing-Review & Editing, Visualization; **Emily Kolenbrander Ho:** Conceptualization, Investigation, Formal analysis, Writing-Review & Editing, Visualization; **Jared E. Toettcher:** Conceptualization, Writing-Review & Editing, Supervision.

## Declaration of competing interests

J.E.T. is a scientific advisor for Prolific Machines and Nereid Therapeutics. The remaining authors declare no conflicts of interest.

## Methods

### *Drosophila* stocks and crosses

Flies were cultured using standard methods at 25°C. *Abd-B-MS2* lines were made in this paper using CRISPR/Cas9 editing described below. Other stocks used include *67;15* (*P{matα-GAL-VP16}67; P{matα-GAL-VP16}15*, Bloomington Drosophila Stock Center (BDSC), #80361), *UAS-optoSOS* (Johnson et al., 2017), y^1^w^118^ (BDSC #6598), *trk*^*1*^ / CyO (BDSC #1212), *tll*-*mCherryLlamaTag* (Ho et al., 2023), *vasa-mCherry* (a gift of Hernan Garcia), *MCP-mNeonGreen* (Singh et al., 2022), and *His2Av-GFP* (a gift of Eric Wieschaus).

For OptoSOS experiments, *67;15* virgins were crossed with *UAS-optoSOS* males. The resulting *67 / UAS-optoSOS; 15 / +* females were used for the experiment. For OptoSOS experiments in a *trk*^*1*^ mutant background *67 trk* ^*1*^*/ CyO; 15* virgins were crossed with *trk*^*1*^*/CyO*; *optoSOS* males to produce *67 trk*^*1*^ */ trk*^*1*^; *15 / UAS-optoSOS* females as in (Johnson et al., 2020).

For live imagining MS2/LlamaTag experiments, a recombinant line was produced containing *MCP-mNeonGreen* and *vasa-mCherry* on Ch2. Virgins from this line were crossed with *tll*-*mCherryLlamaTag* / TM3, Sb males. F1 females containing all three components were then crossed with the desired MS2 males to produce embryos containing all components of the MS2/MCP and LlamaTag systems.

For *trk*^*1*^ mutant experiments (with no OptoSOS), homozygous *trk*^*1*^ females were used.

For all experiments, a cage of parental flies was placed in the dark at room temperature for at least 2 days prior to collection with standard apple juice plates and yeast paste. Except in MS2/MCP experiments, females of the desired maternal genotype were placed in the cage with either males of the same genotype or *y*^*1*^*w*^*118*^ males. For MS2 experiments, males in the cage contained MS2. *His2Av-GFP* flies were used as wild type controls in some experiments.

### Optogenetic stimulation

Embryos were collected on apple juice plates and then stimulated under an array of blue LED lights to induce OptoSOS activity. For long stimulation experiments (2 hours), embryos were collected for 2 hours in the dark, then illuminated for 2 hours under a moderate light intensity of 0.077 mW/cm^2^, and fixed immediately. For short stimulation experiments (5-10 minutes), embryos were collected for 30 min, kept in the dark for 1.5 hrs, illuminated for 5-10 minutes at a higher light intensity of 0.617 mW/cm^2^, returned to the dark for 1-1.5 hrs, and then fixed. These illumination schedules were chosen with the goal of illuminating embryos in late NC13/early NC14 and fixing them at late NC14. In all cases, embryos in the “dark” condition were kept in the dark for the same total duration before fixation. For experiments in which later timepoints were measured (Fig. 1), embryos were returned to the dark for 4-6 hours before fixation (germband extension stage) or 24 hours before cuticle prep.

### Fixation and staining

Embryos were bleached, fixed in an 8% paraformaldehyde/heptane interface, and devitellinized with methanol as described previously (Coppey et al., 2008). HCR v3.0 RNA *in situ* staining was performed according to previous reports (Choi et al., 2018) using initiators and buffers from Molecular Instruments. Probes for *Abd-B* were designed using NM_001316543 (1..1872) which corresponds to the first exon of *Abd-Bm* and an intron on *Abd-Br*. Specific probes for *Abd-Br* were designed using NT_033777.3 (16936105..16943252) which corresponds to an intron of *Abd-Br*. Probes for *tll* (NM_079857.4), *gt* (NM_080310.3), *Kr* (NM_001274252.1), *kni* (NM_079463.3), and *hb* (NM_169233.2) were either ordered from Molecular Instruments or designed using insitu_probe_generator software (Kuehn et al., 2022) and synthesized as an oligo pool by IDT. DAPI was added in one of the final wash steps to stain nuclei.

For Abd-B immunostaining, embryos were rehydrated in PBS + 0.1% Triton X-100 (PBST) and blocked with 20% Western Blocking Reagent (Roche) in PBST (20% WBR). Embryos were treated with primary Mouse anti-Abd-B supernatant (1A2E9 was deposited to the DSHB by Celniker, S.) at 1:10 in 20% WBR overnight at 4°C. After washing, embryos were treated with secondary Goat anti Mouse 647 (Invitrogen) at 1:500 and DAPI for nuclear staining in 20% WBR for 2 hours at room temperature. Following final washes, embryos were mounted using Aqua-Poly/Mount (Polysciences).

Fixed and stained embryos at the desired stage were imaged at 20x on a Nikon A1R or AXR scanning confocal microscope by taking a z stack from the surface to the medial plane.

### Plasmid generation

Guide sequences for MS2 insertion were designed using the FlyCRISPR Optimal Target Finder or taken from published work (Gratz et al., 2014). Sites for MS2 addition were chosen within the 5’UTR of each isoform at a distance from the promoter. The guide sequences (with PAM sequence underlined) were: Abd-B CGACTGCCGAGACATCGTGG<u>GGG</u> and Abd-Br GGTCTGCACCGCTGCTGAAT<u>GGG</u>. Oligos containing the guides were annealed and ligated using T4 ligase into BbsI digested pU6-BbsI-chiRNA vector (Addgene #45946) (Gratz et al., 2013) (**Supplementary Table 1**).

To produce the homologous repair construct for MS2 loop insertion, homology arms extending ∼1 kB in each direction from the target cleavage sites were amplified from genomic DNA that had been extracted from *nos-Cas9* flies (BDSC #78781) (**Supplementary Table 1**) using HiFi polymerase (Takara Biosciences). Homology arms were designed such that the guide sequence would be destroyed by a successful insertion. To facilitate cloning into the pHD-DsRed-24xMS2 donor vector plasmid (a gift of Mike Levine), primers for the left homology arm were flanked by the NdeI restriction site sequence and primers for the right homology arm were flanked by the SpeI sequence. The pHD-DsRed-24xMS2 plasmid was digested with both NdeI and SpeI and the insert and backbone were isolated by gel electrophoresis and purification (NucleoSpin columns, Takara Biosciences). Fragments were assembled using InFusion assembly and transformed into Stellar competent *E. coli* cells (Takara Biosciences). Plasmids were purified using a Qiagen miniprep kit and validated using whole plasmid sequencing (Plasmidsaurus).

### *Drosophila* CRISPR/Cas9 editing

The homology arms and gRNA plasmids were co-injected into nos-Cas9 flies by BestGene Inc and screened for 3xP3-DsRed insertion. Candidate lines were genotyped by PCR with primers flanking the homology arms (**Supplementary Table 1**), and then the linear amplicons were fully sequenced to confirm successful editing (Plasmidsaurus). Sequencing revealed that the Abd-B (R) MS2 line had additional MS2 repeats (28x) but this was considered acceptable. The loxP-flanked 3xP3-DsRed was then removed by crossed to Sp / Cyo-CreW; Dr / TM3 (derived from BDSC #1092) and flies were sequenced again to confirm successful removal.

### Live microscopy

Embryos were mounted for imaging between a semi-permeable membrane and a coverslip in a 3:1 mix of halocarbon 700/27 oil. Live imaging was done at room temperature using a Nikon Eclipse Ti-2 microscope with a Crest V3 spinning disk confocal system at 40x magnification. Imaging started at the beginning of NC14 and images were taken every 1 minute for 1-1.5 hours until ∼30 minutes into gastrulation. Z slices with 1 μm step size were taken to capture the entire nuclear volume of the cell layer.

### Cuticle preparation

Embryos were bleached to dechorionate, mounted on a glass slide in a drop of Hoyer’s Medium, secured with a coverslip, and then heated overnight at 60 °C. Brightfield images were taken with an OptixCam microscope camera. Proportion of embryos with posterior spiracles was counted manually.

### HCR image analysis

All images were processed and analyzed using FIJI (Schindelin et al., 2012). Maximum intensity projections were taken from Z-stack images of each embryo. A line with a width of ∼20 μm was drawn across each embryo, spanning the anterior-posterior axis from the head to the tail. The intensities along the line were then averaged into 100 bins to normalize for embryo length. Then, intensity values were normalized from 0 to 100 within each embryo by subtracting the channel’s minimum intensity value and calculating each position as a percentage of the maximum value. In embryos with no detectable pattern, the minimum value was determined by the averaged background level and the maximum value was set as 3 times this value. Importantly, this intensity normalization within each embryo was necessary because intensity varied between replicates and multiple microscopes were used over the course of this project. Therefore, we do not make absolute claims about signal intensity between conditions, only about patterns within embryos.

### Abd-B immunostaining image analysis

To quantify Abd-B immunostained embryos, a line with a width of ∼20 μm was drawn across each embryo, spanning the anterior-posterior axis. Intensity was measured along this line and normalized to the average intensity of the wild-type Abd-B pattern of *His2Av-GFP* embryos on the same slide. Then, intensity was summed across the quarter of the embryo where Abd-B was expected to be present (For wild-type embryos, this was a center region encompassing the germband-extended tail. For *trk*^*1*^ embryos, this was the most posterior region).

### LlamaTag/MS2 live imaging analysis

Maximum intensity projections were created from the Z slices at each timepoint. Images were then background subtracted by rolling ball subtraction using a radius of 70 pixels in all channels. MS2 bursts were manually identified and marked as white points on the LlamaTag channel. A kymograph was produced from this movie.

### Statistical analysis

All statistical analysis was performed in Prism 11 (GraphPad). Details for statistical tests used can be found in the figure legends. Figure legends indicate the number of embryos for each condition (n). In dot plots, each dot is one embryo. All graphs show the mean ± s.e.m. Significance was defined as *P<0.05, **P<0.01, ***P<0.001 and ****P<0.0001. ns indicates no significance.

## References

Birnie, A., Plat, A., Korkmaz, C., Bothma, J.P., 2023. Precisely timed regulation of enhancer activity defines the binary expression pattern of Fushi tarazu in the Drosophila embryo. Current Biology 33, 2839-2850.e7. 10.1016/j.cub.2023.04.005

Bothma, J.P., Garcia, H.G., Esposito, E., Schlissel, G., Gregor, T., Levine, M., 2014. Dynamic regulation of eve stripe 2 expression reveals transcriptional bursts in living Drosophila embryos. Proc. Natl. Acad. Sci. U.S.A. 111, 10598–10603. 10.1073/pnas.1410022111

Bowman, S.K., Deaton, A.M., Domingues, H., Wang, P.I., Sadreyev, R.I., Kingston, R.E., Bender, W., 2014. H3K27 modifications define segmental regulatory domains in the Drosophila bithorax complex. eLife 3, e02833. 10.7554/eLife.02833

Bronner, G., Jackle, H., 1991. Control and function of terminal gap gene activity in the posterior pole region of the Drosophila embryo. Mech Dev 35, 205–11.

Casali, A., Casanova, J., 2001. The spatial control of Torso RTK activation: a C-terminal fragment of the Trunk protein acts as a signal for Torso receptor in the Drosophila embryo. Development 128, 1709–1715. 10.1242/dev.128.9.1709

Casanova, J., 1990. Pattern formation under the control of the terminal system in the Drosophila embryo. Development 110, 621–628. 10.1242/dev.110.2.621

Casanova, J., Struhl, G., 1989. Localized surface activity of torso, a receptor tyrosine kinase, specifies terminal body pattern in Drosophila. Genes Dev. 3, 2025–2038. 10.1101/gad.3.12b.2025

Casares, F., Sanchez-Herrero, E., 1995. Regulation of the infraabdominal regions of the bithorax complex of Drosophila by gap genes. Development 121, 1855–1866. 10.1242/dev.121.6.1855

Castelli-Gair, J., Greig, S., Micklem, G., Akam, M., 1994. Dissecting the temporal requirements for homeotic gene function. Development 120, 1983–1995. 10.1242/dev.120.7.1983

Choi, H.M.T., Schwarzkopf, M., Fornace, M.E., Acharya, A., Artavanis, G., Stegmaier, J., Cunha, A., Pierce, N.A., 2018. Third-generation in situ hybridization chain reaction: multiplexed, quantitative, sensitive, versatile, robust. Development 145, dev165753. 10.1242/dev.165753

Clark, E., Battistara, M., Benton, M.A., 2022. A timer gene network is spatially regulated by the terminal system in the Drosophila embryo. eLife 11, e78902. 10.7554/eLife.78902

Coppey, M., Boettiger, A.N., Berezhkovskii, A.M., Shvartsman, S.Y., 2008. Nuclear Trapping Shapes the Terminal Gradient in the Drosophila Embryo. Current Biology 18, 915–919. 10.1016/j.cub.2008.05.034

Diaz, R.J., Harbecke, R., Singer, J.B., Pignoni, F., Janning, W., Lengyel, J.A., 1996. Graded effect of tailless on posterior gut development: molecular basis of an allelic series of a nuclear receptor gene. Mechanisms of Development 54, 119–130. 10.1016/0925-4773(95)00467-X

Eldon, E.D., Pirrotta, V., 1991. Interactions of the Drosophila gap gene giant with maternal and zygotic pattern-forming genes. Development 111, 367–378. 10.1242/dev.111.2.367

Gao, Y., Ho, E.K., Toettcher, J.E., 2026. Lights up on the embryonic dance: tools and applications of optogenetics in developmental biology. Genes Dev. genesdev;gad.353459.125v1. 10.1101/gad.353459.125

Gratz, S.J., Cummings, A.M., Nguyen, J.N., Hamm, D.C., Donohue, L.K., Harrison, M.M., Wildonger, J., O’Connor-Giles, K.M., 2013. Genome Engineering of Drosophila with the CRISPR RNA-Guided Cas9 Nuclease. Genetics 194, 1029–1035. 10.1534/genetics.113.152710

Gratz, S.J., Ukken, F.P., Rubinstein, C.D., Thiede, G., Donohue, L.K., Cummings, A.M., O’Connor-Giles, K.M., 2014. Highly specific and efficient CRISPR/Cas9-catalyzed homology-directed repair in Drosophila. Genetics 196, 961–971. 10.1534/genetics.113.160713

Harding, K., Levine, M., 1988. Gap genes define the limits of antennapedia and bithorax gene expression during early development in Drosophila. EMBO J 7, 205–214. 10.1002/j.1460-2075.1988.tb02801.x

Ho, E.K., Kim-Yip, R.P., Simpkins, A.G., Farahani, P.E., Oatman, H.R., Posfai, E., Shvartsman, S.Y., Toettcher, J.E., 2025. A live-cell biosensor of in vivo receptor tyrosine kinase activity reveals feedback regulation of a developmental gradient. Cell Reports 44, 115930. 10.1016/j.celrep.2025.115930

Ho, E.K., Oatman, H.R., McFann, S.E., Yang, L., Johnson, H.E., Shvartsman, S.Y., Toettcher, J.E., 2023. Dynamics of an incoherent feedforward loop drive ERK-dependent pattern formation in the early Drosophila embryo. Development 150, dev201818. 10.1242/dev.201818

Hombría, J.C.G., Rivas, M.L., Sotillos, S.O.L., 2009. Genetic control of morphogenesis - Hox induced organogenesis of the posterior spiracles. International Journal of Developmental Biology 53, 1349–1358. 10.1387/ijdb.072421jc

Hu, N., Castelli-Gair, J., 1999. Study of the Posterior Spiracles of Drosophila as a Model to Understand the Genetic and Cellular Mechanisms Controlling Morphogenesis. Developmental Biology 214, 197–210. 10.1006/dbio.1999.9391

Jimenez, G., Guichet, A., Ephrussi, A., Casanova, J., 2000. Relief of gene repression by torso RTK signaling: role of capicua in Drosophila terminal and dorsoventral patterning. Genes Dev 14, 224–31.

Johnson, H.E., Djabrayan, N.J.V., Shvartsman, S.Y., Toettcher, J.E., 2020. Optogenetic Rescue of a Patterning Mutant. Curr Biol 30, 3414–3424 e3. 10.1016/j.cub.2020.06.059

Johnson, H.E., Goyal, Y., Pannucci, N.L., Schüpbach, T., Shvartsman, S.Y., Toettcher, J.E., 2017. The Spatiotemporal Limits of Developmental Erk Signaling. Developmental Cell 40, 185–192. 10.1016/j.devcel.2016.12.002

Johnson, H.E., Toettcher, J.E., 2019. Signaling Dynamics Control Cell Fate in the Early Drosophila Embryo. Dev Cell 48, 361–370 e3. 10.1016/j.devcel.2019.01.009

Keenan, S.E., Avdeeva, M., Yang, L., Alber, D.S., Wieschaus, E.F., Shvartsman, S.Y., 2022. Dynamics of Drosophila endoderm specification. Proceedings of the National Academy of Sciences 119, e2112892119. 10.1073/pnas.2112892119

Keenan, S.E., Blythe, S.A., Marmion, R.A., Djabrayan, N.J.-V., Wieschaus, E.F., Shvartsman, S.Y., 2020. Rapid Dynamics of Signal-Dependent Transcriptional Repression by Capicua. Developmental Cell 52, 794-801.e4. 10.1016/j.devcel.2020.02.004

Kicheva, A., Briscoe, J., 2023. Control of Tissue Development by Morphogens. Annu. Rev. Cell Dev. Biol. 39, 91–121. 10.1146/annurev-cellbio-020823-011522

Kraut, R., Levine, M., 1991. Spatial regulation of the gap gene giant during Drosophila development. Development 111, 601–609. 10.1242/dev.111.2.601

Kuehn, E., Clausen, D.S., Null, R.W., Metzger, B.M., Willis, A.D., Özpolat, B.D., 2022. Segment number threshold determines juvenile onset of germline cluster expansion in Platynereis dumerilii. Journal of Experimental Zoology Part B: Molecular and Developmental Evolution 338, 225–240. 10.1002/jez.b.23100

Kuziora, M.A., 1993. Abdominal-B protein isoforms exhibit distinct cuticular transformations and regulatory activities when ectopically expressed in Drosophila embryos. Mechanisms of Development 42, 125–137. 10.1016/0925-4773(93)90002-F

Kuziora, M.A., McGinnis, W., 1988. Different transcripts of the Drosophila Abd-B gene correlate with distinct genetic sub-functions. The EMBO Journal 7, 3233–3244. 10.1002/j.1460-2075.1988.tb03190.x

Li, W.X., 2005. Functions and mechanisms of receptor tyrosine kinase Torso signaling: Lessons from Drosophila embryonic terminal development. Developmental Dynamics 232, 656– 672. 10.1002/dvdy.20295

Lu, X., Chou, T.B., Williams, N.G., Roberts, T., Perrimon, N., 1993. Control of cell fate determination by p21ras/Ras1, an essential component of torso signaling in Drosophila. Genes Dev 7, 621–632. 10.1101/gad.7.4.621

Maeda, R.K., Karch, F., 2006. The ABC of the BX-C: the bithorax complex explained. Development 133, 1413–1422. 10.1242/dev.02323

Masuda, L.H.P., Sabino, A.U., Reinitz, J., Ramos, A.F., Machado-Lima, A., Andrioli, L.P., 2024. Global repression by tailless during segmentation. Developmental Biology 505, 11–23. 10.1016/j.ydbio.2023.09.014

Moniot-Perron, L., Moindrot, B., Manceau, L., Edouard, J., Jaszczyszyn, Y., Gilardi-Hebenstreit, P., Hernandez, C., Bloyer, S., Noordermeer, D., 2023. The Drosophila Fab-7 boundary modulates Abd-B gene activity by guiding an inversion of collinear chromatin organization and alternate promoter use. Cell Reports 42, 111967. 10.1016/j.celrep.2022.111967

Morán, É., Jiménez, G., 2006. The Tailless Nuclear Receptor Acts as a Dedicated Repressor in the Early Drosophila Embryo. Mol Cell Biol 26, 3446–3454. 10.1128/MCB.26.9.3446-3454.2006

Pagella, P., Söderholm, S., Nordin, A., Zambanini, G., Ghezzi, V., Jauregi-Miguel, A., Cantù, C., 2023. The time-resolved genomic impact of Wnt/β-catenin signaling. Cell Systems 14, 563-581.e7. 10.1016/j.cels.2023.06.004

Reinitz, J., Levine, M., 1990. Control of the initiation of homeotic gene expression by the gap genes giant and tailless in Drosophila. Developmental Biology 140, 57–72. 10.1016/0012-1606(90)90053-L

Schindelin, J., Arganda-Carreras, I., Frise, E., Kaynig, V., Longair, M., Pietzsch, T., Preibisch, S., Rueden, C., Saalfeld, S., Schmid, B., Tinevez, J.-Y., White, D.J., Hartenstein, V., Eliceiri, K., Tomancak, P., Cardona, A., 2012. Fiji: an open-source platform for biological-image analysis. Nat Methods 9, 676–682. 10.1038/nmeth.2019

Singh, A.P., Wu, P., Ryabichko, S., Raimundo, J., Swan, M., Wieschaus, E., Gregor, T., Toettcher, J.E., 2022. Optogenetic control of the Bicoid morphogen reveals fast and slow modes of gap gene regulation. Cell Rep 38, 110543. 10.1016/j.celrep.2022.110543

Smits, C.M., Dutta, S., Jain-Sharma, V., Streichan, S.J., Shvartsman, S.Y., 2023. Maintaining symmetry during body axis elongation. Current Biology 33, 3536-3543.e6. 10.1016/j.cub.2023.07.050

Smits, C.M., Shvartsman, S.Y., 2020. The design and logic of terminal patterning in Drosophila. Current Topics in Developmental Biology, Gradients and Tissue Patterning 137, 193– 217. 10.1016/bs.ctdb.2019.11.008

Strecker, T.R., Kongsuwan, K., Lengyel, J.A., Merriam, J.R., 1986. The zygotic mutant tailless affects the anterior and posterior ectodermal regions of the Drosophila embryo. Developmental Biology 113, 64–76. 10.1016/0012-1606(86)90108-9

Strecker, T.R., Merriam, J.R., Lengyel, J.A., 1988. Graded requirement for the zygotic terminal gene, tailless, in the brain and tail region of the Drosophila embryo. Development 102, 721–734. 10.1242/dev.102.4.721

Weigel, D., Jürgens, G., Klingler, M., Jäckle, H., 1990. Two Gap Genes Mediate Maternal Terminal Pattern Information in Drosophila. Science 248, 495–498.

Zhao, J., Perkins, M.L., Norstad, M., Garcia, H.G., 2023. A bistable autoregulatory module in the developing embryo commits cells to binary expression fates. Current Biology 33, 2851-2864.e11. 10.1016/j.cub.2023.06.060

