## Supplemental Materials for "The regulatory logic of a dose-dependent developmental fate decision"

Supplemental Figures S1-S2

Supplemental Figure S1 (Related to Figure 1)
Supplemental Figure S2 (Related to Figure 4)

Supplemental Table S1

Supplemental Table S1 (Related to Materials and Methods)

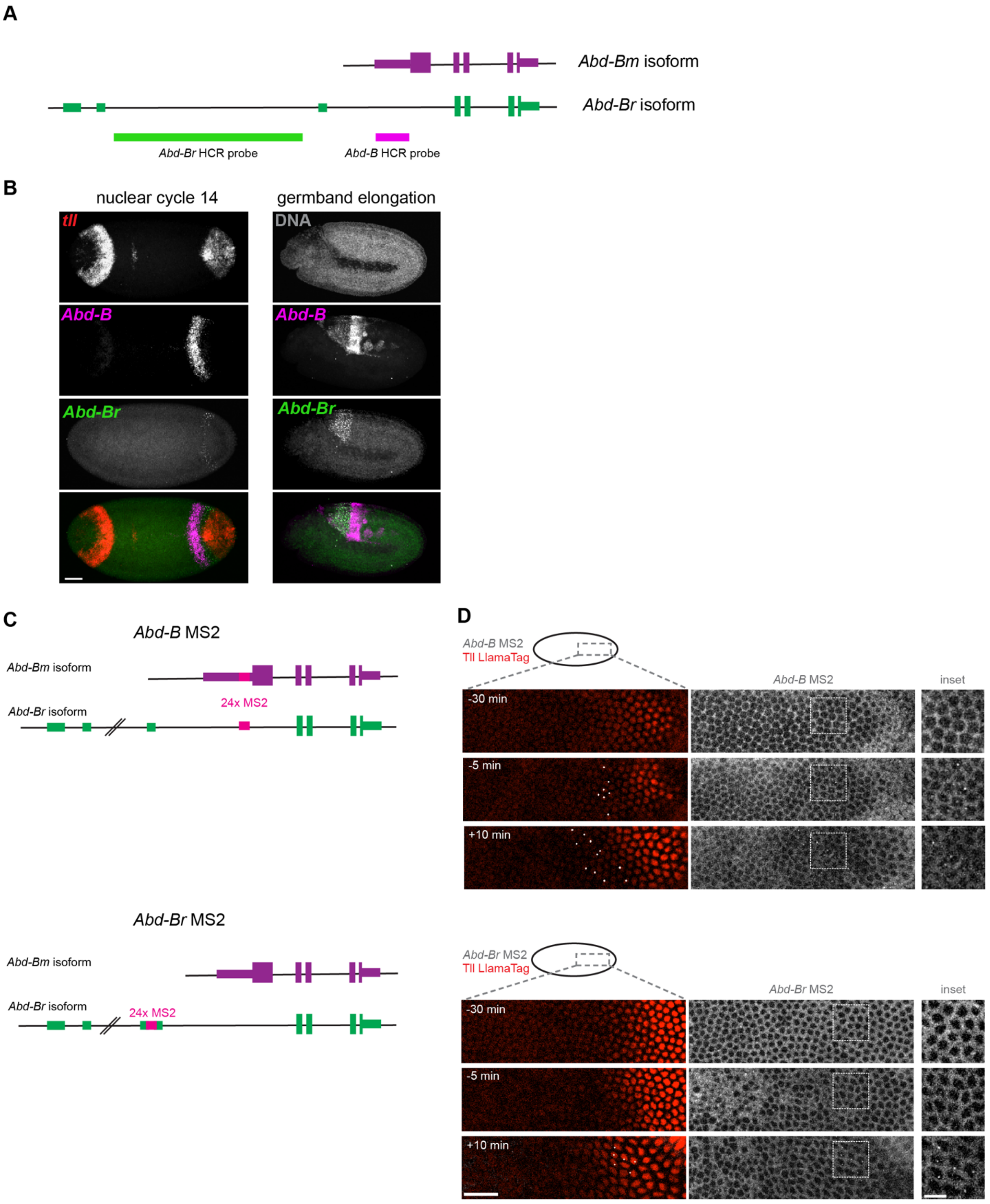


**Supplementary Figure 1: Probes for detecting isoform-specific *Abd-B* expression** (A) Schematic of the two *Abd-B* isoforms: *Abd-Bm* and *Abd-Br*. The Abd-Bm isoform is a longer protein that includes a unique first exon, but a shorter RNA. The Abd-Br isoform is a shorter protein but a longer RNA. Locations used to design isoform-specific HCR probes are indicated. The *Abd-B* HCR probe binds to the unprocessed RNA of both isoforms, but only to the cytoplasmic mRNA of *Abd-Bm*. The *Abd-Br* HCR probe specifically recognizes the unprocessed *Abd-Br* RNA. (B) Maximum intensity projections showing the two *Abd-B* HCR probes as well as the *tll* HCR probe in wild-type nuclear cycle 14 and germband elongation stage embryos. Scale bar 50 μm. In the germband extension stage, the ability of the *Abd-B* probe to detect cytoplasmic *Abd-Bm* mRNA and unprocessed, nuclear *Abd-Br* RNA is clear as these two isoforms are expressed in distinct regions. (C) Schematic of the *Abd-B* isoforms in relation to the position of two 24xMS2 insertions. The first, which we call *Abd-B* MS2, is located in the 5'UTR of the *Abd-Bm* isoform and an intron of the *Abd-Br* isoform and thus should detect both isoforms. The second, which we call *Abd-Br* MS2, is located in the 5'UTR of the *Abd-Br* isoform and thus uniquely detects this isoform. (D) Live imaging of *Abd-B* MS2 (top) and *Abd-Br* MS2 (bottom) together with Tll LlamaTag in the embryo posterior region indicated by the schematic. Time = 0 is the start of gastrulation movements. Images show *Abd-B* MS2 foci and Tll LlamaTag expression 30 minutes before, 5 minutes before, and 10 minutes after gastrulation. In the merged image, *Abd-B* MS2 foci have been manually marked with white dots to allow visualization. The unprocessed *Abd-B* MS2 channel is also shown with an inset enlarging the indicated region. The images shown for *Abd-B* MS2 are the same as in Figure 1E. Scale bars 25 μm, inset: 10 μm.

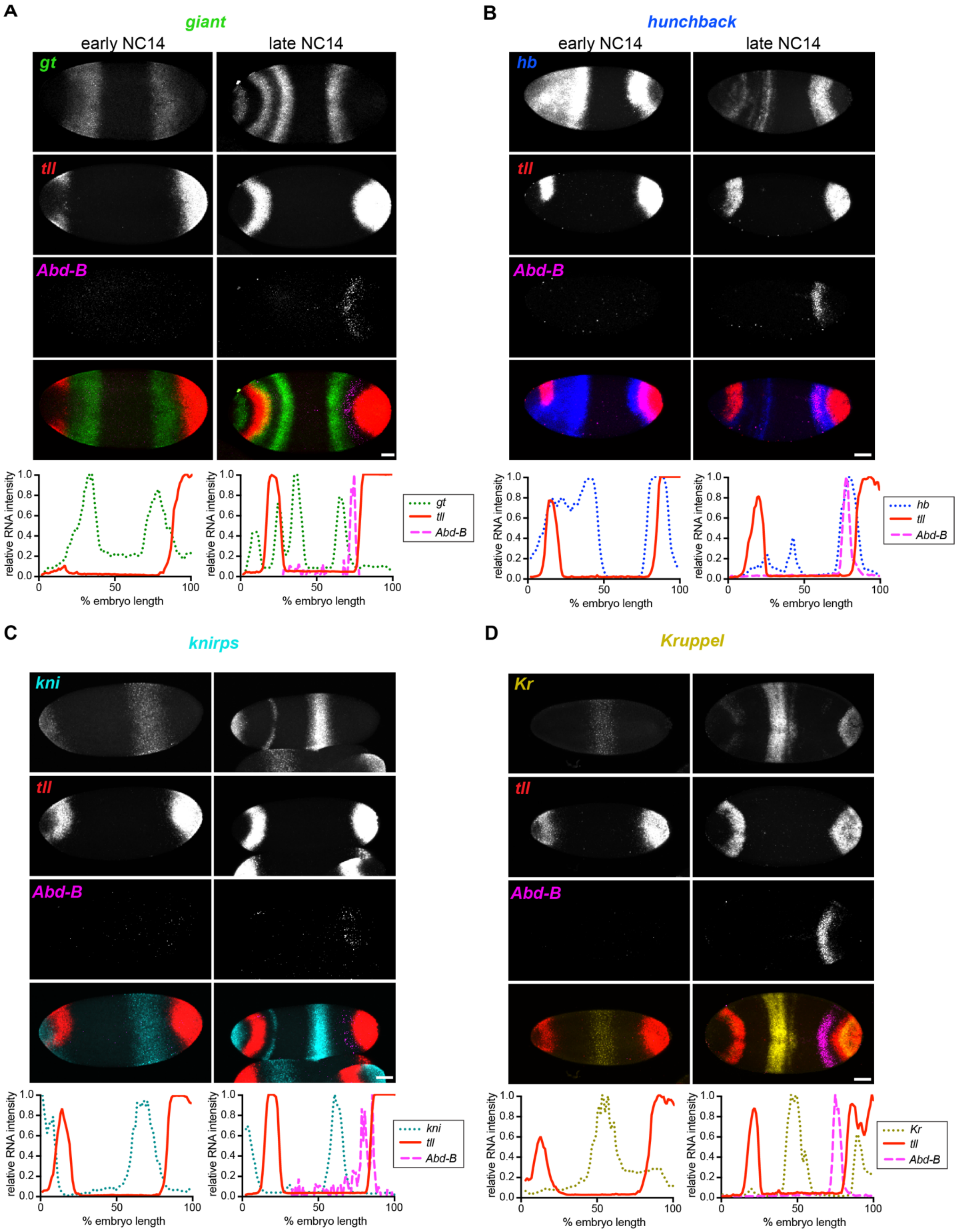


**Supplementary Figure 2: Wild-type expression patterns of gap genes with *Abd-B* and *tll***
Images show early and late stage NC14 embryos stained for *tll*, *Abd-B*, and (A) *gt,* (B) *hb,* (C)

*kni,* or (D) *Kr.* Scale bar 50 µm. Graphs below plot the normalized intensity of each gene across the anterior-posterior axis for the embryo pictured. Abd-B is excluded from the early NC14 plots because it is not expressed at this time. Intensity is normalized from 0 to 100 for each gene and within each embryo.

**Supplementary Table 1.** Primers used for CRISPR/Cas9 and genotyping

| Primer Name | Sequence |
|---|---|
| Abd-B 5' Homology F | GCTAGCGGCCGCGGACATATGGAGGCAAGCGGCCCTGCAACTTC GTCGAGG |
| Abd-B 5' Homology R | ATAAGTACCGTAGTGCATATGTGGGGGTCGTAGAGAGCGCACGCC |
| Abd-B 3' Homology F | AAGTTATAGAAGAGCACTAGTCGATGTCTCGGCAGTCGGGTCGCACGCC |
| Abd-B 3' Homology R | TGCATGGAGATCTTTACTAGTAAGTGGAACGACCCCGTTCCCAATTTAACGTCCCC |
| Abd-B Guide F | CTTCGCGACTGCCGAGACATCGTGG |
| Abd-B Guide R | AAACCCACGATGTCTCGGCAGTCGC |
| Abd-B Genotyping F | CCAGAGCCAGTCCCAGTCGAAGTGCGATACC |
| Abd-B Genotyping R | CGTGGGTGGCGCTGTGGCCATAC |
| Abd-Br 5' Homology F | GCTAGCGGCCGCGGACATATGTGGGTTGGGAATTGGAGCAGGAAGACGTCATTGTGC |
| Abd-Br 5' Homology R | ATAAGTACCGTAGTGCATATGAATGGGGATGGATCTACTCCACCGGTTTGCTCACTTCC |
| Abd-Br 3' Homology F | AAGTTATAGAAGAGCACTAGTCAGCAGCGGTGCAGACCCACCAAGCACC |
| Abd-Br 3' Homology R | TGCATGGAGATCTTTACTAGTATCGCGACCAATTATACGAGTCGTCAAGCGCAGTGAC |
| Abd-Br Guide F | CTTCGGTCTGCACCGCTGCTGAAT |
| Abd-Br Guide R | AAACATTCAGCAGCGGTGCAGACC |
| Abd-Br Genotyping F | GCGCAGCATCTCCGGCCGGAGAACC |
| Abd-Br Genotyping R | GGTATCGCACTTCGACTGGGACTGGCTCTGG |
